# Emergence of rhythmic chunking in complex stepping of mice

**DOI:** 10.1101/2023.03.02.530893

**Authors:** Kojiro Hirokane, Toru Nakamura, Yasuo Kubota, Dan Hu, Takeshi Yagi, Ann M. Graybiel, Takashi Kitsukawa

**Affiliations:** Graduate School of Life Sciences, Ritsumeikan University, Kusatsu, Shiga, Japan; Graduate School of Frontier Biosciences, Osaka University, Suita, Osaka, Japan; McGovern Institute for Brain Research and Dept. of Brain and Cognitive Sciences, Massachusetts Institute of Technology, Cambridge, Massachusetts, United States

## Abstract

Motor chunking is important for motor execution, allowing atomization and efficiency of movement sequences. However, it remains unclear why and how chunks contribute to motor execution. To analyze the structure of naturally occurring chunks, we trained mice to run in a complex series of steps and identified the formation of chunks. We found that intervals (cycle) and the positional relationship between the left and right limbs (phase) of steps inside the chunks, unlike those outside the chunks, were consistent across occurrences. Further, licking by the mice was also more periodic and linked to the specific phases of limb movements within the chunk. Based on these findings, we propose the *rhythm chunking hypothesis*, whereby within chunks, the repetitive movements of many body parts are linked by the rhythm parameters: cycle and phase. The computational complexity of movement may thereby be reduced by adjusting movements as the combination of rhythms.

## Introduction

Many of the movements that we perform in our daily lives are continuous movements in which many body parts move together in coordination (Walshe, 1947). In order for such continuous movements to be successfully performed, these body parts must be coordinated both spatially and temporally. For example, when you are playing the piano, your fingers work together one after another to produce the music. When we learn a continuous movement consisting of a series of motor elements, the movement sequence is initially performed by making each motor element, but after many repetitions, several successive adjacent elements gradually form small clusters, which are called chunks (Gallistel, 1980; Lashley et al., 1951; Willingham, 1998).

A variety of studies has suggested that the positive effects of movement chunking include movement efficiency and energy conservation. One of the behavioral tasks most frequently employed for research on chunking is the discrete sequence production (DSP) task and its derivatives, in which subjects perform a series of key presses (Gallistel, 1980; Paillard, 1960). Experiments based on this task indicate that motor chunks are formed during continuous movement, and that the entire sequence could collectively be generated as a response to a single stimulus. As demonstrated by Verwey *et al*. (1999), the reaction times to start the first tapping of a keypress sequence was shortened through performance of the DSP task, and Yamaguchi and Logan (2014) demonstrated that the time needed to perform the entire sequence decreased with learning. Given that the time required to start the movement sequence becomes shorter, the time to prepare the series of movements may be reduced, indicating the reduction of complexity of information processing in motor chunks as compared to the generation of each movement of the sequence one by one. Ramkumar et al. (2016) analyzed jerk parameters of arm movements in non-human primates using a continuous reaching task to model the chunk as a local optimal control problem. They found that local optimal control allows a counterbalance between efficiency and complexity of movement, so that chunking contributed to the reduction of computational complexity and was cognitively efficient.

Chunks that form naturally in our daily life often involve movements of multiple body parts requiring coordination among these parts. However, most of the chunks studied so far have been repetitions of simple single movements of body parts, and many of them have been movements with specific instructions given to human subjects. In many cases, the boundaries of the chunk, the pace of the repetitive movement, and the movement to be performed were pre-determined in the experimental plans. For this reason, the formation, structure, and function of chunking in continuous movements requiring coordination remain to be further clarified.

Here, we gave mice the opportunity to form chunks spontaneously in the process of learning and performing complex sequential movements (see STAR Methods for details). We examined how chunks are generated, how they grow to include increasing numbers of individual movements, and how movements differ between those inside and outside of the chunks that are formed. For this purpose, we used the step-wheel task (Kitsukawa *et al*., 2011) to train mice to perform complex continuous movements. The step-wheel is a vertical running wheel for mice with pegs for footholds arranged in a ladder-like or complex pattern on the running surface. The wheel turns at a constant speed under computer control, and water-restricted mice are required to run with complex stepping patterns without any specific instructions for the timing of each step in order to obtain reward. We have used this step-wheel task to examine sequential learning and performance of mice, including genetically modified mice (Crittenden et al., 2021; Kitsukawa et al., 2011; Nakamura et al., 2014, 2017, 2020). In this study, we found that as the training on complex steps progresses, the chunked sequences in which the movement parameters were stable became more pronounced and were gradually elongated, and that the motor coordination within the chunks was supported by the rhythm of these movements.

## Results

### Stable running regions in a complex peg pattern

To test how chunks are formed spontaneously out of a series of different movements, it was necessary to train mice first to perform repetitive, yet non-uniform, continuous movements over a certain period of time. For this purpose, we used a training protocol in which the mice repeatedly ran 50–60 times on 12 irregularly placed pegs on each side of the step-wheel. After the initial use of the regular peg-pattern to familiarize them with the wheel, they were exposed, in each session, to 24 pegs (pegs 1 to 24) arranged in the complex peg-pattern (Fig. 1A–D).

**Fig. 1.**
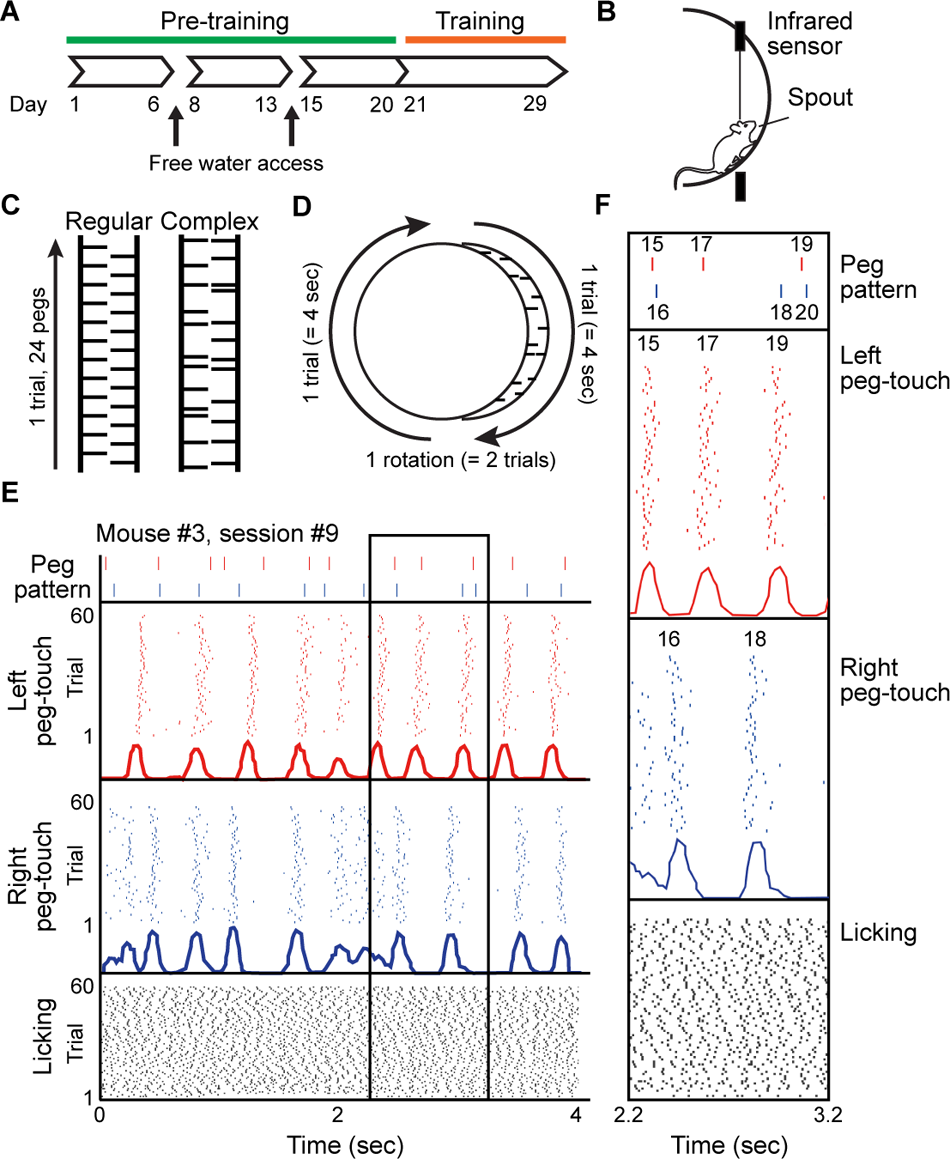
The step-wheel and peg touches of mice. A. Timeline of the entire experiment, including pre-training and training periods. B. Side view of the step-wheel. An infrared beam was equipped to detect a mouse when it was close to the spout. The turning speed of the wheel was kept constant at 12.5 cm/sec in all sessions by computer control. C. Peg patterns used in the study. The Regular peg-pattern was used for pre-training, and the Complex peg-pattern was used for the training. A single pattern consisting of 24 pegs was defined as 1 trial. D. Diagram illustrating the relationship between trials and rotation. Within one rotation of the step-wheel, the same pattern was repeated twice. E. Raster plots and histograms of the timing of left and right peg touches (middle) by a mouse running the Complex peg-pattern (top), and raster plots of the timing of licking (bottom). F. Enlarged view of the raster plots and histograms in the box in E.

A key feature of a motor chunk is that it represents a stable movement pattern (Popp et al., 2020). Thus, we sought to find stable regions in the running of mice in the wheel. In stable regions, the timing of the peg-touches should vary little from trial to trial. On the other hand, unstable regions should show high variability. Therefore, we measured the timing of all the limbs as the paws touched the pegs, and we determined whether the stability of touch timing of the footfalls differed for different pegs. Raster plots and histograms of the peg-touch times (Fig. 1E) illustrate examples indicating that the variability of the peg-touch timing was different depending on the location in the entire peg-pattern This variability suggested that there were stable and unstable regions in the running.

To quantify the variability of the locomotion in the stable and unstable regions during the runs, we first calculated the inter-touch intervals, taken as the time interval between two consecutive peg-touches comprising a single step, and we calculated the coefficient of variation of the inter-touch intervals (inter-touch CVs) for all pegs of each trial (Fig. 2A). It was apparent without quantification that the inter-touch CVs decreased as training progressed. In fact, measurements showed that the mean inter-touch CVs of all mice decreased across sessions, and when the values of session 1 were compared to those of each other session, the values were significantly lower after session 4 (Fig. 2B, black line, n = 7, p = 2.4 × 10^−6^ by one-factor ANOVA; p < 0.05 by Tukey-Kramer test). The smaller inter-touch CV at the later stages of the training is consistent with our previous results reporting the acquisition of peg-patterns on the step-wheel (Nakamura et al., 2017).

**Fig. 2.**
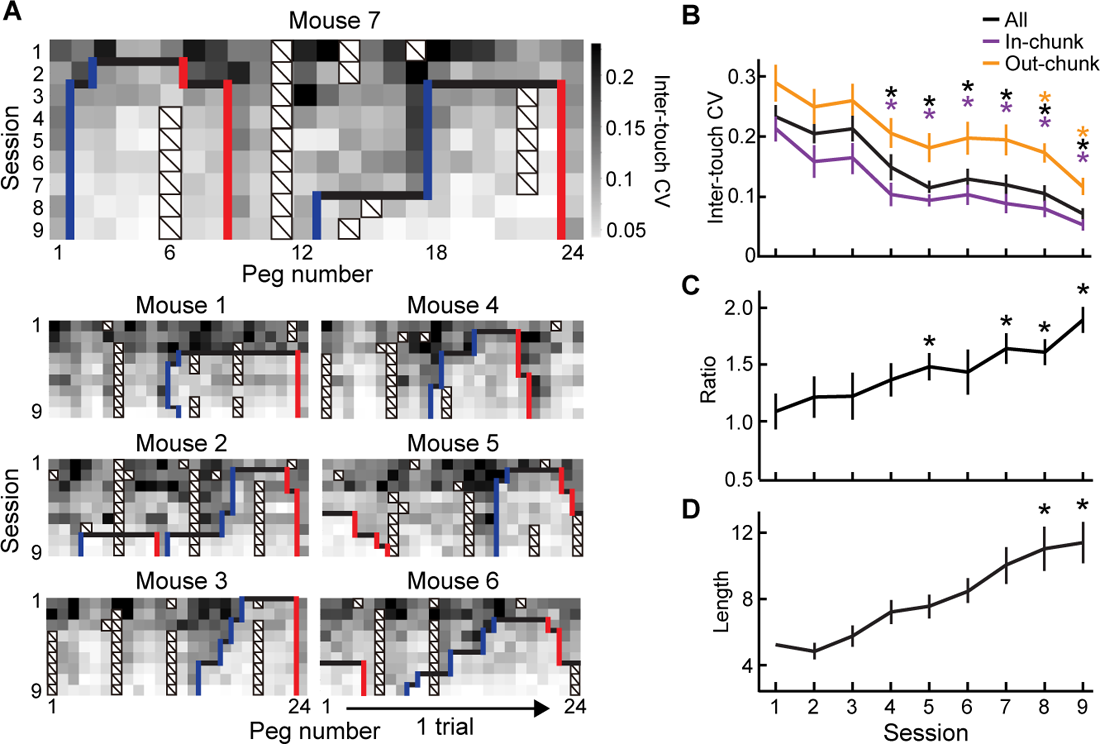
Emergence of motor chunks during running on the step-wheel detected by the inter-touch CV. A. Heat maps of the inter-touch CV between consecutive peg pairs in all sessions (session1–9, vertical axis) for all mice (n = 7). The start (blue line) and end (red line) of the chunk are shown. Specifically, the inter-touch CV at the n^th^ peg is the coefficient of variation of the time interval from the peg-touch to the (n − 1)^th^ peg to that of the n^th^ peg within one session. The pegs that were excluded from the analysis due to the low number of touches in a session are marked with diagonal lines. One trial (= 24 pegs) is shown as the horizontal axis. B. The inter-touch CV (mean ± SEM, n = 7) for all pegs (black), in-chunk pegs (purple) and out-chunk pegs (orange) through the training. Black, purple and orange asterisks indicate, respectively, the significant decrease in the inter-touch CV compared to that of session 1 for all (p = 2.4 × 10^−6^, by one factor ANOVA; *p < 0.05, by Tukey-Kramer test), in-chunk (p = 4.5 × 10^−4^, by one factor ANOVA; *p < 0.05, by Tukey-Kramer test) and out-chunk (p = 1.2 × 10^−3^, by one factor ANOVA; *p < 0.05, by Tukey-Kramer test) pegs. C. The ratio of the inter-touch CV between in-chunk and out-chunk pegs is shown as the mean ± SEM (n = 7, p =1.8 × 10^−3^, by one factor ANOVA; *p < 0.05, by Tukey-Kramer test). D. The average length of chunks in all mice across sessions (n = 7, p = 2.2 × 10^−6^, by one factor ANOVA; *p < 0.05, by Tukey-Kramer test).

However, there were high-CV segments and low-CV segments even in later sessions (Fig. 2A). This finding indicated there were regions in which touch intervals varied from trial to trial within a session as the mice ran on the pegs with high inter-touch CVs. At the pegs with low inter-touch CVs, the mice ran with similar inter-touch intervals during most trials. This meant that the steps within low-CV segments were stably repeatable, suggesting that the continuous movements in these segments might comprise chunks.

If they were chunks, we expected that (1) there would be a low-CV segment with roughly constant inter-touch CVs; (2) there would be unstable segments with high inter-touch CVs before and after the low-CV segment; and (3) the inter-touch CVs would be different inside and outside of a low-CV segment. Therefore, we identified low-CV segments, which were considered chunks, by several criteria (see Supplementary Fig.1 and STAR Methods for details). We call the first and last pegs of the low-CV segments as, respectively, the start-peg and end-peg. Examples of the start-peg and end-peg of the chunks identified by this method are indicated, respectively, by blue and red vertical lines in Fig. 2A. Here, the total lengths (transverse) displayed in Fig. 2A represent single lengths of peg-patterns (trials).

We were then able to determine whether the decreases in inter-touch CV occurred inside the chunks or outside of the chunks, or across both regions. In other words, was the stable part further improved, or was there an improvement in the unstable part? To answer this question, we separately calculated the inter-touch CVs of touches inside the chunk (in-chunk) and outside the chunk (out-chunk). We found that both in-chunk and out-chunk inter-touch CV decreased over the course of training. The in-chunk inter-touch CVs in session 1 were significantly higher than those after session 4 (Fig. 2B, purple, n = 7, p = 4.5 × 10^−4^ by one-factor ANOVA; p < 0.05 by Tukey-Kramer test). The out-chunk inter-touch CV also decreased with the progress of training, with the average inter-touch CV in session 1 being significantly higher than those after session 8 (Fig. 2B, orange, n = 7, p = 1.2 × 10^−3^ by one-factor ANOVA: p < 0.05 by Tukey-Kramer test). Throughout the 9 sessions of training, the average inter-touch CVs were always smaller for in-chunk, followed by those for total session and then out-chunk inter-touch CVs. In our previous study (Nakamura et al., 2017), it was reported that each mouse reduced the variance of its steps overall with training, but without providing a definition of in-chunk and out-chunk periods. In the current study, we conducted the same experiment with different mice, and we performed the same analysis, comparing data for inside and outside of the chunk.

To estimate the relative reduction in the inter-touch CV between out-chunk and in-chunk periods, we calculated the ratio of in-chunk to out-chunk inter-touch CVs. We found significant increases in the ratio in sessions 5, 7, 8, and 9 compared to the ratio in the first session (Fig. 2C, n = 7, p = 1.8 × 10^−3^ by one-factor ANOVA; p < 0.05 by Tukey-Kramer test). Thus, the out-chunk inter-touch CVs were higher than the in-chunk inter-touch CVs especially in the later sessions, indicating that the chunked regions of the runs gradually became demarcated by lower CV as training proceeded. This pattern seems reasonable, given that the in-chunk and out-chunk inter-touch CVs decreased with roughly similar slopes and the in-chunk inter-touch CVs were always smaller than the out-chunk CVs, leading to larger ratios in the late stages of the training. This result supported the validity of the method that we developed here to set the chunk boundaries.

These results appear to indicate that the chunk locations became more fixed in the later sessions. However, contrary to this impression, the chunk boundaries changed from session to session (Fig. 2A). The changes were always in the direction of elongating the length of chunked sequence; the start-pegs moved backward, and the end-pegs moved forward with respect to the direction of running (Fig. 2A, blue and red lines, respectively). We found that there was a significant increase in absolute chunk length in sessions 8 and 9 compared to the length in session 2 (Fig. 2D, n = 7, p = 2.2 × 10^−6^ by one-factor ANOVA; p < 0.05 by Tukey-Kramer test). This result suggests that the increase in the CV ratio across learning does not necessarily correspond to a fixation of chunk boundaries. Rather, chunk boundaries were found to be changing dynamically during the course of training. Further, we found no concatenation of small chunks. The development of chunks may be caused by gradual elongation at the two ends of a given chunk rather than by concatenation.

### Individual chunking strategies reflected in the stepping patterns of the mice

We next addressed the issue of the locations of chunks within the runs on the complex peg-pattern. They appeared relatively similar in a given individual, but different across different individuals (Fig. 2A). To compare their locations, we defined a chunk vector (Supplementary Fig. 2) with 24 components corresponding to 24 pegs, each assigned a value of either 1 (inside chunk) or 0 (outside chunk). The similarities of chunk locations between sessions of the same mouse (intra-individual) or of different mice (inter-individual) were calculated by the cosine similarities of the chunk vectors. The matrix showing the cosine similarities of chunk vectors of all sessions of all mice indicated that they tended to be higher across sessions of the same mouse (diagonal blocks) and lower across sessions of different mice (non-diagonal blocks, Fig. 3A). The averages of the cosine similarities between the intra-individuals and the inter-individuals were significantly different (Fig. 3B, n = 7, p = 6.4 × 10^−20^ by two-tailed t-test). This result indicates that the chunk locations were relatively constant within a given mouse but that different individuals formed chunks at different locations even with the identical complex peg-pattern. This finding implies that the chunks that we identified were developed independently by individual mice, rather than simply being dependent on the peg pattern.

**Fig. 3.**
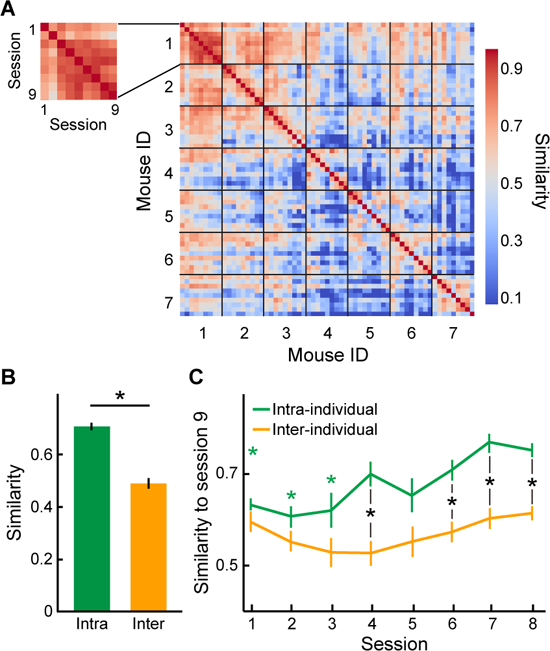
Development of chunks through successive sessions and individual differences in these chunk formation behaviors. A. Heat map of products of chunk vectors, showing similarity between sessions within individual mice (e.g., inset) and between sessions from different mice. B. Intra- and inter-individual comparison of chunk similarity (mean ± SEM). The cosine similarities of the chunk vectors were averaged for intra-individual (green) and inter-individual (orange) sessions (*p = 6.4 × 10^−20^, by two-tailed t-test). C. Chunk similarities between each of sessions 1–8 to the ninth session. Chunk similarities of intra-individual (green) and inter-individual (orange) sessions are displayed as the mean ± SEM. Green asterisks indicate the sessions whose intra-individual chunk similarities to session 9 are significantly low compared to that of session 8 (p = 6.9 ×10^−6^, by one factor ANOVA; *p < 0.05, by Tukey-Kramer test). No significance detected for inter-individual comparison. Black asterisks indicate the sessions in which the intra-individual similarities are significantly higher compared to the inter-individual similarities of the same session (*p = 5.6 × 10^−10^ on session 4; *p = 8.4 × 10^−6^ on session 6; *p = 5.5 × 10^−10^ on session 7; *p = 9.1 × 10^−11^ on session 8, by two-tailed t-test).

To estimate the developmental progression of chunk locations, that is, whether the differences started from the beginning of training or gradually increased, we compared the cosine similarities of the chunk vectors of sessions 1–8 to that of the last session (session 9). We found that the cosine similarities of the intra-individual chunk vectors became gradually larger, indicating that the chunking pattern over successive sessions became more similar to that of the final session. When compared to the similarity between sessions 8 and 9, those between sessions 1–3 and 9 were significantly smaller (Fig. 3C, green, n = 7, p = 6.9× 10^−6^ by one-factor ANOVA; p < 0.05 by Tukey-Kramer test). This finding was important in demonstrating that chunk formation was not stable already in the early sessions, but instead, gradually, as training proceeded, became similar to those of the final session. We found no such experience-related increase in similarity in the comparisons of training sessions from different individuals (Fig. 3C, orange, n = 7, p = 0.14 by one-factor ANOVA). Furthermore, the similarity score was higher for the intra-individual comparisons than the inter-individual comparisons in the sessions 4, 6, 7 and 8 (Fig. 3C, black asterisks, n = 7, two-tailed t-test; p = 5.6×10^−10^ in session 4; p = 8.4×10^−6^ in session 6; p = 5.5×10^−10^ in session 7; p = 9.1×10^−11^ in session 8). These results indicate that a running pattern specific for each individual might have been acquired by the middle phase of training, and that, thereafter, the chunks became more elongated based on that running strategy of each individual mouse, contrary to the idea that the running strategies converged to one particular running pattern determined by the peg pattern.

### Limb interval preserved inside the chunk

It is an open question how movements are structured inside of chunks and how movements differ inside and outside the chunks. To address this question, we explored the difference of movements made inside and outside of chunks, with a specific focus on the coordination of right and left forelimb movements, measured by the timing of the footfalls recorded and concentrating on the coordination of the forelimb movements rather than on individual footfalls. We used two indices of coordination: the footfall *intervals* as an index of coordination of repetitive movements of a single limb (Fig. 4A), and the positional relationship between the left and right legs (*phase*) as an index of coordination between the left and right limbs (Fig. 4B).

**Fig. 4.**
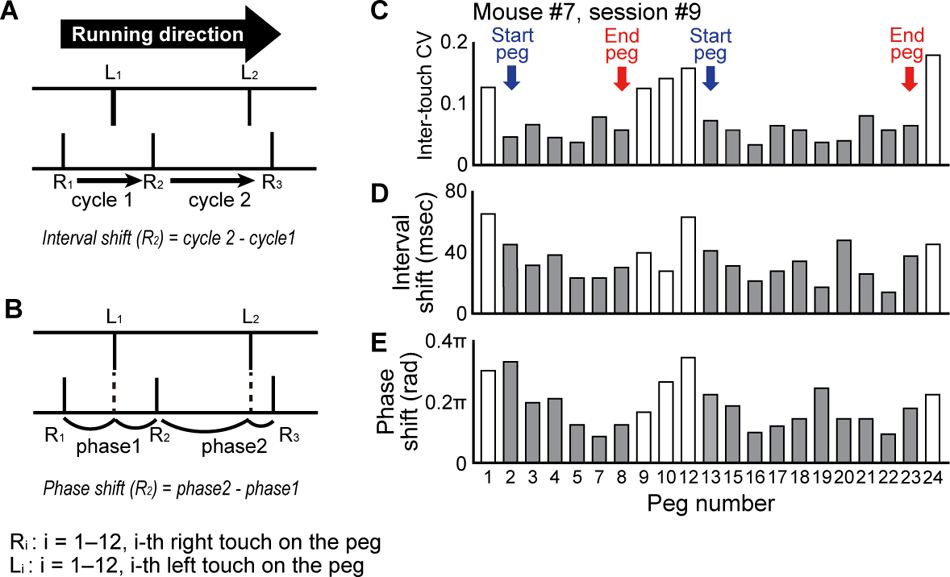
Calculations and examples of inter-touch CV, interval-shift and phase-shift. A. Scheme for describing the formulas for calculating the interval-shift. *R_i_* indicate the *i*-th touch to the right side peg. We calculated the interval-shift in the following calculation: 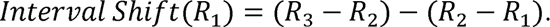 B. Scheme for describing the formulas for calculating the phase-shift. *L_i_* and *R_i_* indicate, respectively, the *i*-th touch to the left and right pegs. We calculated the phase-shift as: 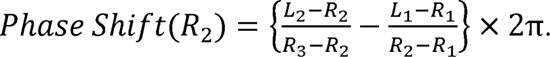 C–E. An example of inter-touch CV (C), interval-shift (D), and phase-shift (E) of a session performed by mouse number 7 on day 9. Start-pegs, end-pegs and chunk region (gray) are shown in C.

In continuous locomotion, individual body parts are used repeatedly, so that whether or not the intervals between two consecutive repetitive movements change from one repetition to the next should have a profound impact on the continuous locomotion. Such changes as occur with repetition can be considered as representing coordination on a temporal axis between consecutive movements of the same body part. To test whether such coordination emerge during the step-wheel training, we calculated, for the left and right forelimbs, the *interval-shift,* which is the difference between successive inter-touch intervals. In other words, interval-shift is an index of how much change there was in the time taken for one cycle before and one cycle after a given peg of interest. The inter-touch intervals were distributed around an average of 369.73 ± 0.55 msec (Fig. 5A). The interval-shifts were distributed around an average of 41.53 ± 0.13 msec (Fig. 5B).

**Fig. 5.**
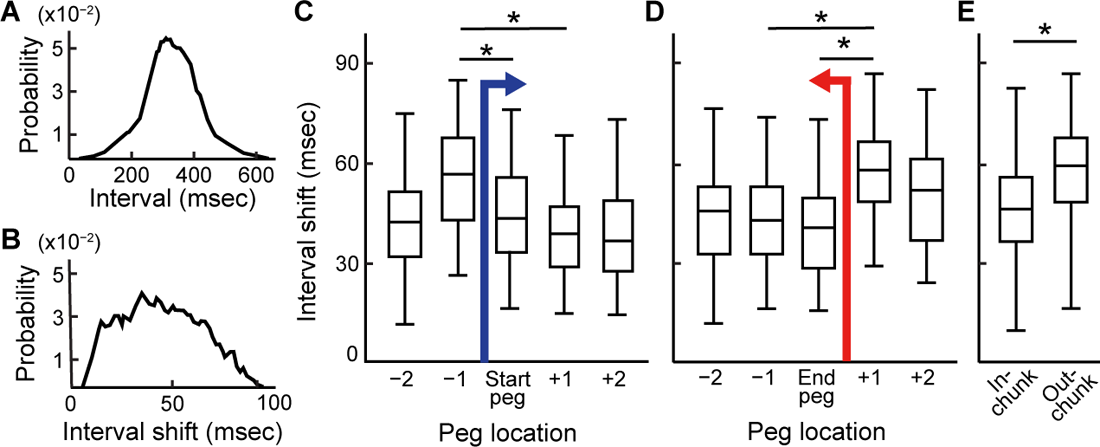
The modulation of interval-shifts around the chunk boundaries. A, B. Distribution of intervals of steps (A) and interval-shift (B) in all sessions of all mice. C, D. Box plots of interval-shift aligned at start-peg (C; p = 7.8×10^−3^, by one factor ANOVA; *p < 0.05, by Tukey-Kramer test) and at end-peg (D; p = 4.1×10^−3^, by one factor ANOVA; *p < 0.05, by Tukey-Kramer test) for all chunks from all session, including both right and left pegs. The median and interquartile range are shown. E, Comparison of the interval-shifts between in-chunk and out-chunk periods (*p = 5.6 × 10^−6^, by two-tailed t-test). The median and interquartile range are shown in boxplots.

We found that the interval-shift was high where the inter-touch CV was high, that is, adjacent to the start-pegs and end-pegs (Fig. 4C, D). To clarify the positional relationship between a putative chunk and the interval-shift, we aligned the interval-shift to the start-peg or end-peg (Fig. 5C, D). We used the start-pegs and end-pegs after session 4 (Fig. 3C), when chunk formation had become stable. This analysis showed that the interval-shifts at the start-peg and the next peg were significantly smaller than those at the peg before the start-peg (Fig. 5C, p = 7.8×10^−3^ by one-factor ANOVA; p < 0.05 by Tukey-Kramer test). The interval-shifts at the end-peg and the peg before the end-peg were also significantly smaller than those of the peg after the end-peg (Fig. 5D, p = 4.1×10^−3^ by one-factor ANOVA; p < 0.05 by Tukey-Kramer test). These results indicated that the changes in time taken for consecutive cycle were smaller within the chunk than outside the chunk.

To quantify this relationship, we compared the interval-shifts between in-chunk and out-chunk periods. The average of the in-chunk interval-shifts was significantly smaller than that of the out-chunk interval-shift (Fig. 5E, p = 5.6 × 10^−6^ by two-tailed t-test). These results clearly demonstrated that the time durations for successive cycle were stable inside the chunk, indicating that the touch intervals in the chunk segment were relatively constant compared to touch intervals outside of the chunk. We next tested whether the modulation of the interval-shift around the chunk boundary differed among individual mice. The results demonstrated that, across all the mice, a similar trend occurred, with differences in significance level of the modulation in interval-shift among the mice (Supplementary Fig. 3).

### Inter-limb phase preserved inside the chunk

We defined the interval-shift on a per-limb basis to demonstrate that the cycles of each body part were modulated around the chunk region, but the stepping of the animal is determined by a combination of the left and right limb movements. Therefore, we next focused on the phase of the limb movements, which we defined as the time difference between left and right paw touches in a cycle. The phase was distributed around an average of 0.971 ± 2.32 × 10^−3^ (Fig. 6A, radian), indicating that the mice prefer to move left and right limbs alternately rather than simultaneously. The phase-shift, the difference of the phase between two consecutive cycles, showed a skewed distribution with a thicker tail to the right (Fig. 6B, mean ± SEM: 0.220 ± 1.12 × 10^−3^ rad; median, 0.184 π rad). The phase-shifts were often observed to be high where the inter-to^π^uch CV and the interval-shifts were high, that is, neighboring the start-pegs and the end-pegs (Fig. 4C–E). To investigate the relationship between the chunk position and the phase-shift, we aligned the phase-shift to the start-peg or end-peg (Fig. 6C, D), and found that the phase-shifts at the start-peg and the next peg were significantly higher than that at the second peg before and after the start-peg (Fig. 6C, p = 6.9×10^−3^ by one-factor ANOVA; p < 0.05 by Tukey-Kramer test). When compared to the phase-shift of the end-peg, the phase-shift of two pegs before and after the end-pegs was significantly higher (Fig. 6D, p = 1.6 × 10^−4^ by one-factor ANOVA; p < 0.05 by Tukey-Kramer test).

**Fig. 6.**
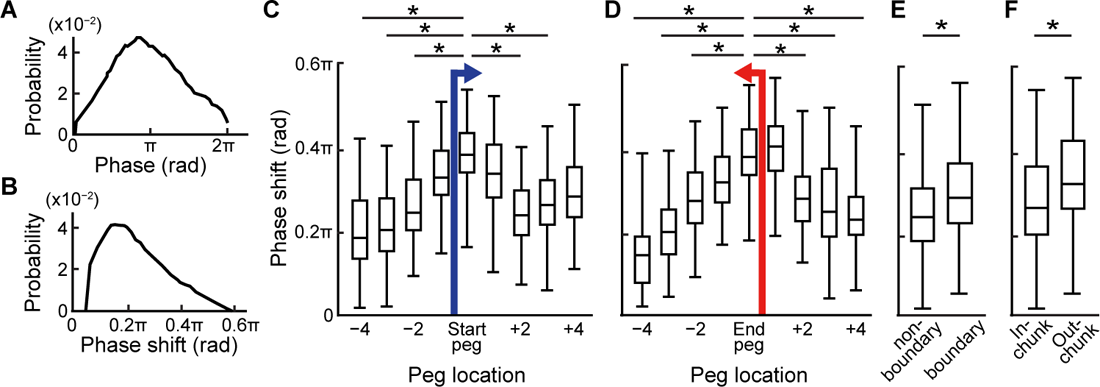
The modulation of phase-shifts at chunk boundaries. A, B. Distribution of phase in a step (A) and phase-shift (B) in all sessions of all mice. C, D. Box plots of phase-shift aligned by the start-peg (C; p = 6.9×10^−3^, by one-factor ANOVA; *p < 0.05, by Tukey-Kramer test, compared to the start-peg) and by the end-peg (D; p = 1.6×10^−4^, by one-factor ANOVA; *p < 0.05, by Tukey-Kramer test, compared to the end-peg) for all chunks from all session. The median and interquartile range are shown in boxplots. E, F. Comparison of phase-shifts (mean ± SEM) between the chunk boundary segment and the non-boundary segment (E; *p = 7.2 × 10^−18^, by two-tailed t-test) and between inside and outside the chunk (F; *p = 5.5 × 10^−6^, by two-tailed t-test,) for all chunks from all session. The median and interquartile range are shown in boxplots.

These results indicated that the phase-shifts might be higher at the boundaries of chunk regions, unlike the interval-shifts, in which a significant difference was observed between inside and outside the chunk. Therefore, we compared the phase-shifts between the boundaries, which were defined as three consecutive pegs centered at the start-peg or end-peg, and the other regions in the peg-pattern (non-boundary regions). The phase-shift was significantly higher in the boundary regions (Fig. 6E, p = 7.2 × 10^−18^ by two-tailed t-test). Even when the phase-shift was compared between in-chunk and out-chunk periods as we did in the interval analysis, a significant difference was observed (Fig. 6F, p = 5.5 × 10^−6^ by two-tailed t-test). Additionally, we examined the modulation of the phase-shift in relation to the chunk boundary on a per-mouse basis and found a similar trend across all the mice, with differences in significance level of modulation in phase-shift among the mice (Supplementary Fig. 4).

These findings indicate that the phase in the chunk regions was relatively constant compared to the phase in parts of the runs outside the chunk. This result in turn implies that the coordination of left and right limbs was relatively consistent within the chunk, as compared to their relative coordination during non-chunked parts of the runs. The phase-shift increased around the chunk boundaries (Fig. 6), whereas the interval-shift increased just outside the chunk (Fig. 5). This finding indicates that the adjustment of step intervals may have been made before the adjustment of step phases at the start of the chunk, whereas the adjustment of step phase may have been made before the adjustment of step intervals at the end of the chunk. For both situations, the chunk proved a critical variable in the adjustments.

### Stable relationship between licking and touch inside the chunk

The fact that the coordination of the left and right forelimbs differed between in-chunk and out-chunk periods raised the intriguing question of whether movements of other parts of the body that are not directly involved in running were also coordinated with limb movements. As a first step toward addressing this issue, we detected licking by means of a voltage sensor attached to the spout, and we analyzed the licking patterns registered (Fig. 1E, F). The mice licked to drink water while they were running in the wheel, and their licking was quite periodic, with a mean interval of 117.5 ± 0.22 msec (Fig. 7A, B). We examined the relationship between licking and peg-touches inside and outside chunks and found that the timing of licking was aligned with the timing of peg-touches for each peg (Fig. 7C). The licking was periodic with about 100 msec peak-to-peak latency when aligned to some pegs (Fig. 7C, top), but not when aligned to other pegs (Fig. 7C, bottom), indicating that the coordination of licking and forelimb movement was different for each peg.

**Fig. 7.**
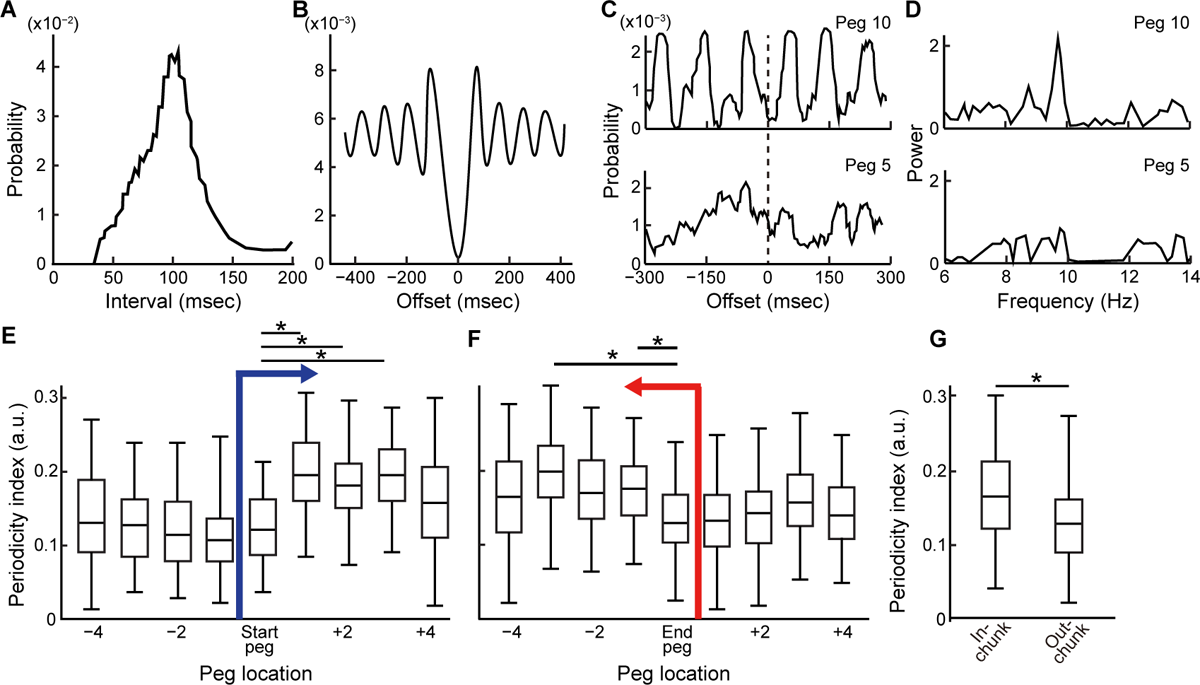
The modulation of licking periodicity around chunk boundaries. A. Distribution of the lick intervals. B. Auto-correlogram of lick timing. C. Cross-correlograms of licks aligned to the time of peg 10 (top) and peg 5 (bottom) touches. Note that the cross-correlogram to peg 10 shows large amplitude whereas that to peg 5 shows small amplitude. The data used to construct the cross-correlogram were from session #9 of mouse #6. D. Power spectrums calculated from the cross-correlograms shown in C. Note that the power spectrum for peg 10 (top) shows a large peak at around 9 Hz, which corresponds to licking frequency, whereas power spectrum for peg 6 (bottom) does not. E, F. Box plot of periodicity indices for each peg aligned to the start-peg (E; p = 3.4×10^−3^, by one factor ANOVA; *p < 0.05, by Tukey-Kramer test) and to the end-peg (F; p = 6.1×10^−3^, by one factor ANOVA; *p < 0.05, by Tukey-Kramer test) for all chunks from all session. The median and interquartile range are shown in boxplots. G. Comparison of periodicity indices between in-chunk and out-chunk pegs (*p = 2.1 × 10^−8^, by two-tailed t-test) for all chunks from all session. The median and interquartile range are shown in boxplots.

To identify pegs that were accompanied by periodic licking, we calculated the power spectrum of the cross correlogram that aligned the licks with respect to touches to the pegs (Fig. 7D). We defined a licking periodicity index for each peg as the maximum value of the power spectrum in the range of 8–10 Hz, which corresponds to licking frequency observed. Thus, the higher this index value, the more periodically the mice licked the spout around the peg. To investigate the relationship between the periodicity index and the locomotor chunks, the periodicity index of each peg was aligned with the start-peg and the end-peg. We found that the periodicity index increased significantly at 1–3 steps after the start-peg compared to that of the start-peg (Fig. 7E, p = 3.4×10^−3^ by one-factor ANOVA; p < 0.05 by Tukey-Kramer test), and that the periodicity index significantly increased 1–3 steps before the end-peg compared to that of the end-peg (Fig. 7F, p = 6.1×10^−3^ by one-factor ANOVA; p < 0.05 by Tukey-Kramer test). This pattern suggested that the coordination of forelimb movement and licking was higher inside of the chunk than outside of the chunk. Thus, we compared the periodicity index between in-chunk and out-chunk periods. We found that the periodicity index was significantly higher inside the chunk than outside of it (Fig. 7G, p = 2.1 × 10^−7^ by two-tailed t-test). It was important to test these results also for each mouse. We did so (Supplementary Fig. 5) and found that the mice exhibited individual differences in significance level of modulation in licking periodicity, but we found that there was a similar trend in the results across all mice.

Because there was a possibility that the licking itself could hardly be performed during the out-chunk periods, the auto-correlogram of the licking in each region (Supplementary Fig. 6A) and the average drinking frequency in each region (Supplementary Fig. 6B) were calculated. We confirmed that licking occurred rhythmically even in the out-chunk period, although it was slightly less rhythmic than during in-chunk periods (Supplementary Fig. 6A). In addition, the number of licks was quite similar between in-chunk (6.46 ± 1.37/sec) and out-chunk (6.22 ± 1.25/sec; Supplementary Fig. 6C. These results were important in indicating that the licking in the chunk was more periodically linked to peg-touches than it was outside of the chunks. Inside the chunk, the interval and phase of steps were not only stable and highly coordinated, but also the licking, which was also periodic, was coordinated with the limb movements.

## Discussion

Here we developed a protocol to examine the formation of stable running patterns by mice as they navigated a complex peg-pattern of footholds (pegs) that they were required to run on in order to receive reward delivered during the runs. Three major findings emerged. First, as the mice learned the patterns, they developed stable, repetitive several step stepping of their runs, regions that we defined a chunk. The intervals of cycles were more consistent in these chunks than outside of the chunks, suggesting that the movements of limbs was iso-periodic and cycles were more rhythmic within the chunk. Second, we observed that the relative positions of the left and right forelimb touches comprising a full cycle were stable within the chunks, indicating that the rhythmic movements of each forelimb were coordinated with phase, which is a parameter of rhythm. This further indicates that within the chunked parts of the runs, the coordination of movements among body parts was achieved through links between rhythms via phases. Third, we found that the rhythm of licking, which occurred at a different frequency than the stepping and was not directly related to running, was phase-locked with step rhythms especially within chunks, indicating that in chunked regions, the movements of many body parts can be coordinated with each other by virtue of the rhythm.

According to Thompson et al., 2019, there are three commonly accepted definitions of chunks: (1) the number of chunks should increase, (2) the proportion of actions falling within a chunk should increase, and (3) chunks should be faster than non-chunks. In this study, the turning speed of the wheel was kept constant and thus the running speed of mice was constant. This procedure allowed us for a detailed comparison of inside and outside the chunks, but it became impossible to satisfy the definition of (3). Therefore, instead of definition (3), speed increase, we used the reproducibility of step timing in the definition of chunks. In tasks in which the animal can change the speed of movement by themselves, the movement in chunks should be faster, but in tasks where the animal cannot change the speed, such as running in the step-wheels used in this study, we considered that the movement in the chunk region would be more highly reproducible instead of being faster, because of a speed-accuracy trade-off relationship (Fitts’ law, Fitts, 1954). Indeed, it has been found in both monkey and rat studies that when a behavior is sufficiently repeated, that behavior becomes stabilized (e.g., Barnes et al., 2005; Desrochers et al., 2010). The details of the movements after the behavior were stabilized were not measured in these studies. But we did analyze these in detail, and we found that the stabilized movements in mouse stepping was based on rhythm.

Based on these findings, we propose the *rhythm chunking hypothesis* whereby the motor chunk is composed of the combination of rhythms generated by the repetition of simultaneously ongoing movements of multiple body parts (Fig. 8). To generate optimal movements involving multiple body parts, it should be necessary to consider the number of possible combinations of all joints and muscles to be adjusted. In addition, in continuous movements, the degrees of freedom for the timing of such combinations are infinite. However, by using rhythm as the coordinating parameter—matching the cycles of each movement, connecting them in a particular phase, and repeating them—the degrees of freedom can be greatly reduced (see Graybiel & Grafton, 2015). This should also save brain resources used for movement generation.

**Fig. 8.**
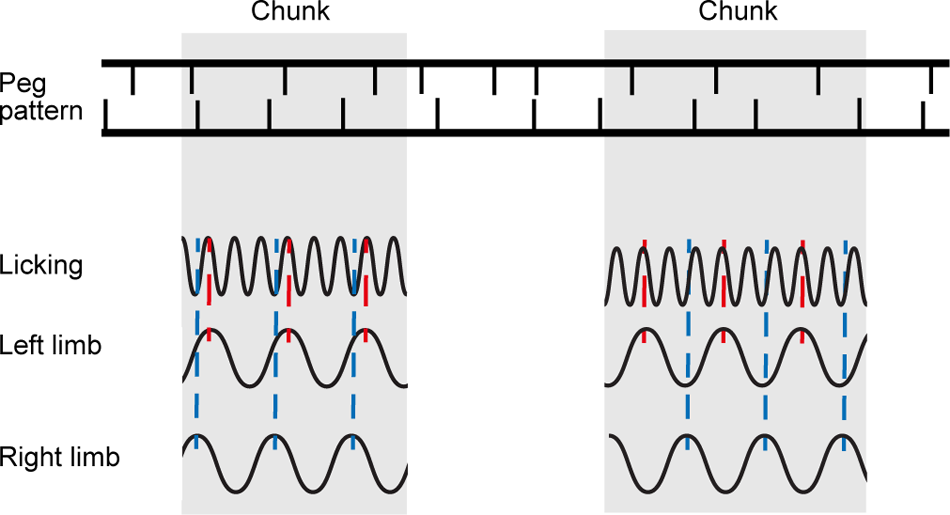
Schematic diagram illustrating the occurrence of motor chunks structured by rhythmic coordination of licking, left forelimb and right forelimb. Motor chunks are composed of the combination of rhythms for stepping and licking of a mouse running in the step-wheel.

The positions of the body parts determined by rhythm coordination will often not be the optimal position for each individual movement. In the case of stepping on the complex peg-pattern, the optimization of movements for each step may be impaired to some extent. However, if the best positions of the body parts for each step are selected, extra efforts should be needed for the transition between steps, which may not be optimal for continuous stepping. If the positional relationship of the body parts in a consecutive step is constrained to be within a certain range, the movements formed by the rhythm coordination could be smoother and more efficient in the transition. As a result, even if the movement at each step were not optimal, continuous movements as a whole might become more efficient, which might be the driving force to form the chunk. The longer the movement sequence generated by the rhythmic coordination, the more efficient and advantageous. This efficiency could be the determinant of chunk length.

The application of the rhythm chunking strategy may be limited to repetitive movements. However, most of the continuous movements that we perform routinely can be regarded as repetitive movements. For example, we need to move our arm forward and then back when we walk, and we need to close our mouth and then reopen it when we talk. Furthermore, some discrete movements like throwing a ball can be considered as a cycle of repetitive movements. The rhythmic chunking of movements thus can be applied to a wide range of movements. It is unclear whether the motor chunks previously reported in earlier studies were rhythm chunks, but the chunks reported using periodic movements may well be rhythm chunks (Ruitenberg et al., 2013; Verwey & Abrahamse, 2012).

It is known that rhythms are often combined in simple integer ratios automatically in continuous movements, which implicates that the coordination of periodic movements occurs via links of rhythm parameters: interval (cycle) and phase. For example, when an animal walks or runs, the rhythms of gait and breathing are combined in a simple ratio. In the gallop of quadrupeds, the ratio is 1:1 (Alexander McN., 1989; Bramble & Carrier, 1983). Even in the bipedalism of humans, the rhythm of walking and breathing has a relationship of simple frequency ratios, such as 2:1 and 3:1 (Bernasconi & Kohl, 1993; Persegol et al., 1991). In addition, whisking rhythms were also shown to be synchronized with sniffing in the rat with simple ratios such as 2:1, 1:1 and 1:2 (Ranade et al., 2013). The existence of many of such examples indicates that rhythm coordination is a general phenomenon for motor coordination.

In this study, we observed how mice developed complex continuous movements by examining their behavior on the step-wheel, and we found that small chunks were generated, and that the length of the chunks increased as training proceeded. Because the movements were coordinated in the chunk by the parameters of rhythm, the increase in chunk lengths indicates that the regions in the peg pattern that mice can continue to perform without major changes requiring the rhythm parameters to become longer. In addition, the fact that the difference in inter-touch CV between inside and outside of the chunks increased as the training progressed suggests the possibility that the magnitude of differences in the rhythm parameters may increase at chunk boundaries in the later sessions. We did not observe chunk concatenation, which might have resulted from difficulty in merging chunks with different rhythm parameters.

We found individual differences among mice in the location of chunks within the peg patterns that they ran. This observation made it clear that the location of chunk formation was not determined solely by the peg pattern, given that all mice ran on the same peg pattern; and it provided evidence against the possibility that the chunk was formed at easy-to-run regions determined by the peg pattern. Once formed, chunks tended to be maintained in successive sessions, and individual differences increased as training progressed. Thus, the running movements may be fixed to a different movement pattern for each mouse, rather than converging on the most optimal movement physically determined by the peg pattern.

The different convergence patterns for individual mice may depend on the position of the first chunk that the mouse formed early in training. In complex peg-patterns, there can be multiple strategies for executing successful movements consisting of multiple steps. Each mouse may have selected one of them, probably by chance, and continued to use that strategy thereafter. The diversity of individual chunking strategies observed in our experiments is consistent with the results of a previous study (Popp et al., 2020), in which different tapping sequences formed across individual subjects. Alternatively, differences in the features that consist of chunks may have affected where the chunks form. At the boundaries of the detected chunks, both of the rhythm parameters, interval and phase, changed, but the magnitude of the changes varied among individuals. It is possible that the location of chunk formation differed depending on whether the emphasis was on interval or phase. It has been reported that the importance of features in a chunk differs when there is a difference in the position at which the chunk is formed (Takeya et al., 2021). It may be true for chunks in general that the features that consist of a chunk affect the location of chunks. As well, the differences among mice may have resulted from differences in sensory conditions. Mice could run in dark conditions with the same proficiency as they did under light conditions. However, when their whiskers were trimmed, they were unable to run proficiently in the wheel (Crittenden *et al*., 2021). These findings suggest that the differences in whisker sensibility could be involved in the individual differences in chunk locations.

On the basis of all of these considerations, we propose the hypothesis that motor chunks are developed by and are dependent on the coordination of rhythms of the moving parts. Reconstructing complex repetitive movements into a combination of rhythmic coordination of the relationships among them may reduce the calculations needed to generate effective movement sequences. This may reduce the load on the brain, eliminating the need to be aware of each individual movement, and automating continuous movement, which may lead eventually to the formation of habits (Graybiel, 2008).

### Limitations of the study

We acknowledge that there are several limitations in our study. First, in our experimental system, the complexity of the peg pattern was represented by 12 pegs on each side. With this design, there is no guarantee that all possible complexity would have had appeared within the complex peg-pattern, and a longer peg pattern would have revealed more about the chunks. We found that the length of chunks increased, but they were not concatenated. We consider that this elongation may be the principle of chunk formation, but if the mice were required to learn and perform a longer peg pattern, concatenation might have occurred, as reported in other studies (Verwey, 2001; Wright et al., 2010). Second, we did not analyze hindlimb movements, even though the presence or absence of rhythm coordination between forelimbs and hindlimbs is a very important issue. This lack of data acquisition occurred because touches were detected by voltage sensors located only at the spout, and we could not record both forelimbs and hindlimbs simultaneously. In the future, if both forelimbs and hindlimbs can be simultaneously recorded by high-speed video, it will be possible to determine whether, as we would expect, the movements of all four limbs are coordinated by rhythm.

## Acknowledgements

We thank the members of both laboratories for valuable discussions and help. This work was funded by JST SPRING (JPMJSP2138 to K.H.), KAKENHI (18H04945 and 22K18662 to T.K.), and the National Institute of Mental Health (R01 MH060379 to A.M.G.).

## Author contribution

A.M.G, T.K. designed the experiments. K.H., T.N., Y.K., D.H., and T.K. performed the experiments. K.H., T.N., and T.K. performed the data analysis. K.H., Y.K, A.M.G., and T.K. wrote the paper. T.Y., A.M.G., and T.K. supervised the project.

## Figure legends

**Supplementary Fig. 1.**
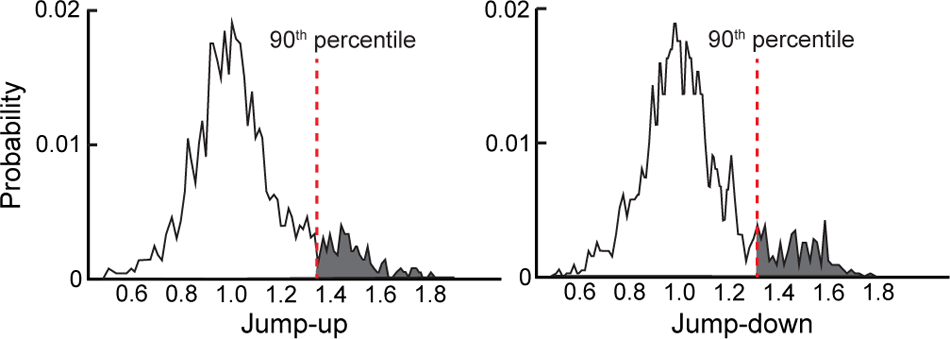
A distribution of jump-ups and jump-downs. Distributions of jump-ups and jump-downs of inter-touch CV were calculated across all sessions for all mice. Upper 10 percentile values are indicated by red dotted lines, and upper 10 percentile of the distributions are indicated by shading.

**Supplementary Fig. 2.**
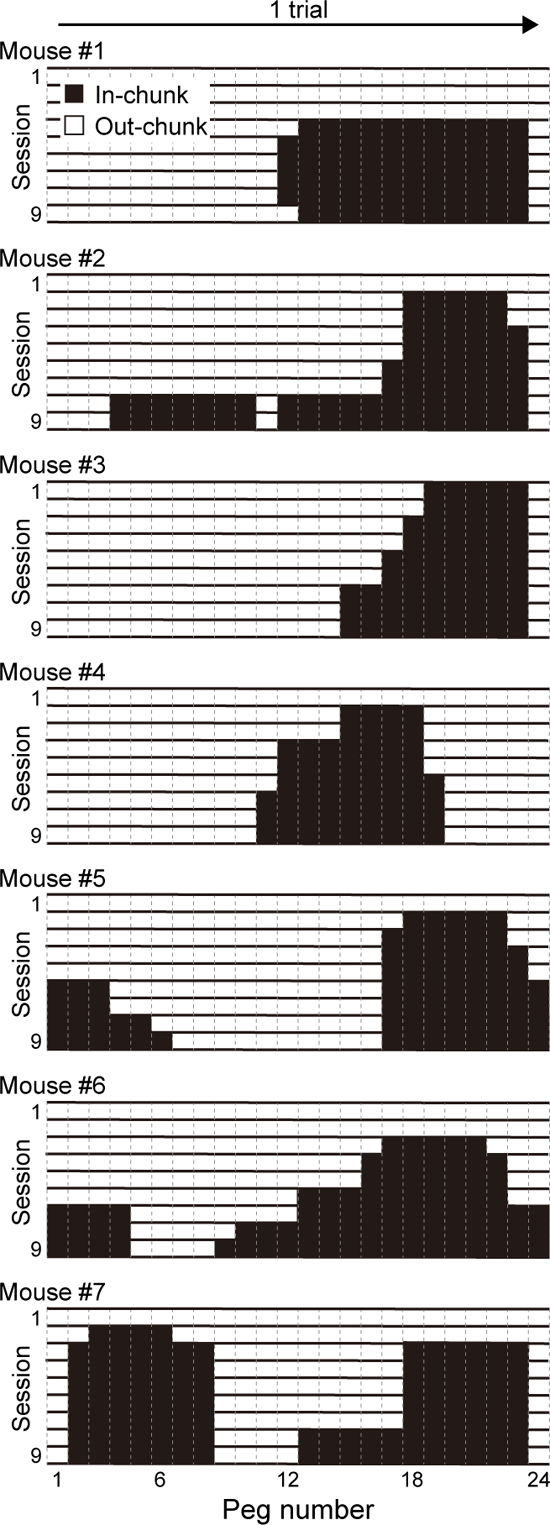
Chunk locations represented by the binary map of chunk vectors. Chunk vectors shown in the binary map for all mice (n = 7) and all sessions (sessions 1–9). The chunk vector comprised of as many elements as pegs in a pattern (24). Thus, the size of the chunk vector is (1, 24). The black regions represent the regions identified as a chunk.

**Supplementary Fig. 3.**
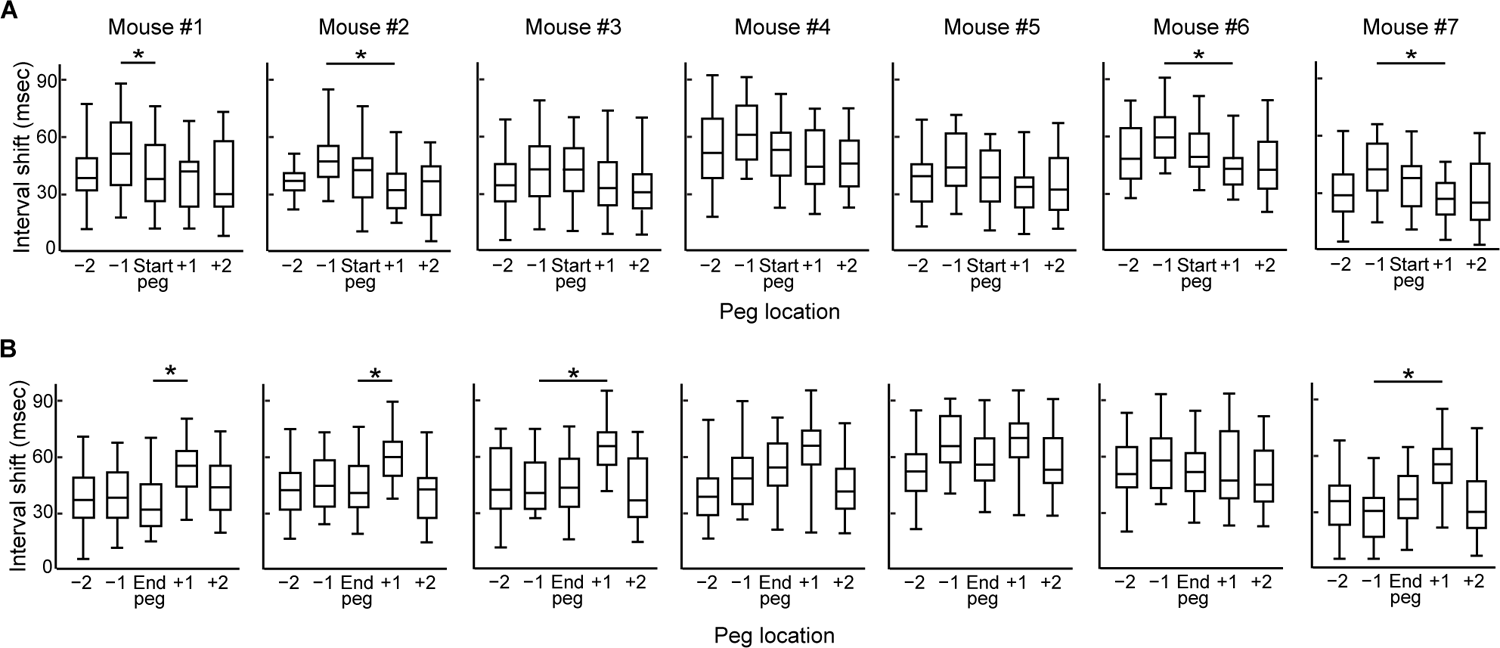
Modulation of interval-shift around the chunk boundary for each mouse. A. Box plots of interval-shift aligned to the start-peg for each mouse (Mouse #1: p = 6.2×10^−3^, Mouse #2: p = 5.1×10^−5^, Mouse #3: p = 0.061, Mouse #4: p = 0.25, Mouse #5: p = 0.083, Mouse #6: p = 2.3×10^−2^, Mouse #7: p = 3.1×10^−2^, by one factor ANOVA; *p < 0.05, by Tukey-Kramer test). The median and interquartile range are shown. B. Box plots of interval-shift aligned to the end-peg for each mouse (Mouse #1: p = 2.9×10^−4^, Mouse #2: p = 2.1×10^−2^, Mouse #3: p = 8.2×10^−4^, Mouse #4: p = 0.21, Mouse #5: p = 0.31, Mouse #6: p = 0.092, Mouse #7: p = 5.3×10^−3^, by one factor ANOVA; *p < 0.05, by Tukey-Kramer test). The median and interquartile range are shown.

**Supplementary Fig. 4.**
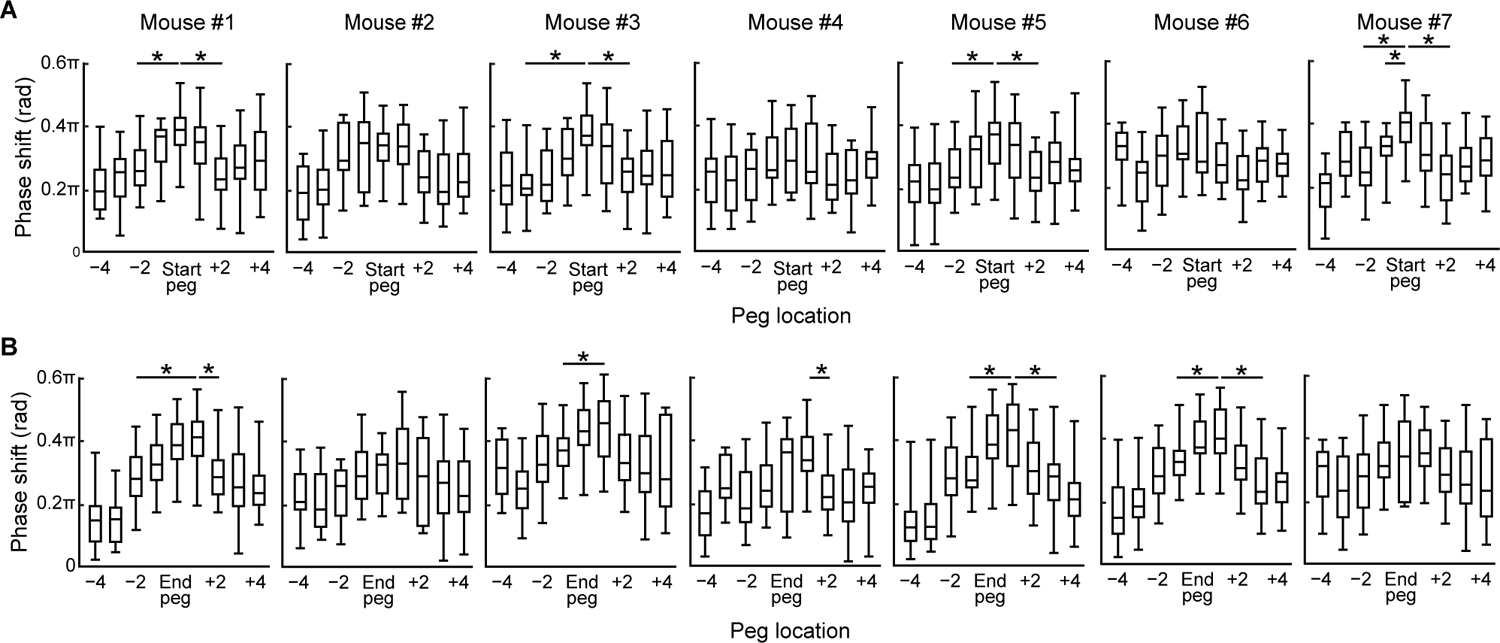
Modulation of phase-shift around the chunk boundary for each mouse. A. Box plots of phase-shift aligned to the start-peg for each mouse (Mouse #1: p = 3.2×10^−5^, Mouse #2: p = 6.2×10^−4^, Mouse #3: p = 5.2×10^−3^, Mouse #4: p = 0.42, Mouse #5: p = 9.1×10^−4^, Mouse #6: p = 0.35, Mouse #7: p = 1.5×10^−5^, by one factor ANOVA; *p < 0.05, by Tukey-Kramer test). The median and interquartile range are shown. B. Box plots of phase-shift aligned to the end-peg for each mouse (Mouse #1: p = 1.8×10^−4^, Mouse #2: p = 0.41, Mouse #3: p = 1.2×10^−3^, Mouse #4: p = 6.2×10^−6^, Mouse #5: p = 5.6×10^−3^, Mouse #6: p = 7.2×10^−5^, Mouse #7: p = 5.2×10^−2^, by one factor ANOVA; *p < 0.05, by Tukey-Kramer test). The median and interquartile range are shown.

**Supplementary Fig. 5.**
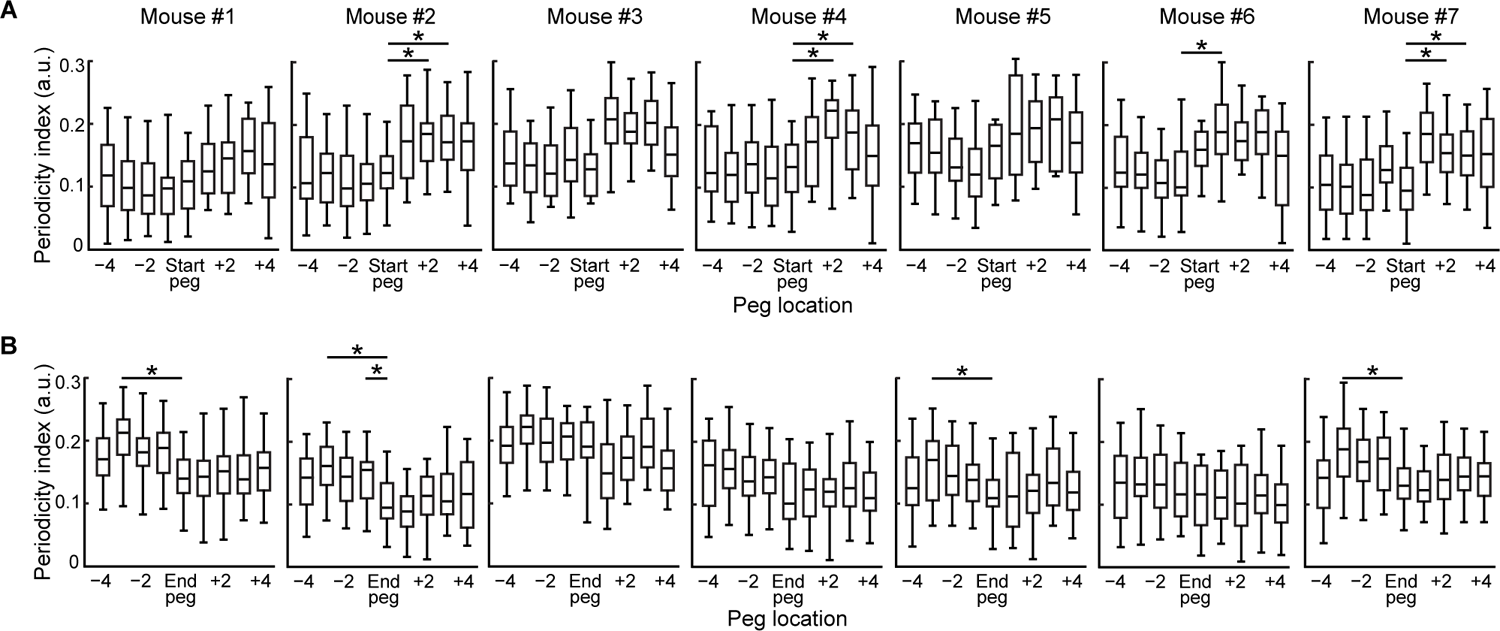
Modulation of licking periodicity around the chunk boundary for each mouse. A. Box plot of periodicity indices for each peg aligned to the start-peg for each mouse (Mouse #1: p = 0.32, Mouse #2: p = 1.3×10^−4^, Mouse #3: p = 2.5×10^−3^, Mouse #4: p = 1.1×10^−4^, Mouse #5: p =0.36, Mouse #6: p = 1.9×10^−2^, Mouse #7: p = 3.5×10^−6^, by one factor ANOVA; *p < 0.05, by Tukey-Kramer test). The median and interquartile range are shown. B. Box plot of periodicity indices for each peg aligned to the end-peg for each mouse (Mouse #1: p = 7.8×10^−4^, Mouse #2: p = 9.3×10^−4^, Mouse #3: p = 5.0×10^−3^, Mouse #4: p = 0.082, Mouse #5: p = 0.59, Mouse #6: p = 0.41, Mouse #7: p = 6.9×10^−4^, by one factor ANOVA; *p < 0.05, by Tukey-Kramer test). The median and interquartile range are shown.

**Supplementary Fig. 6.**
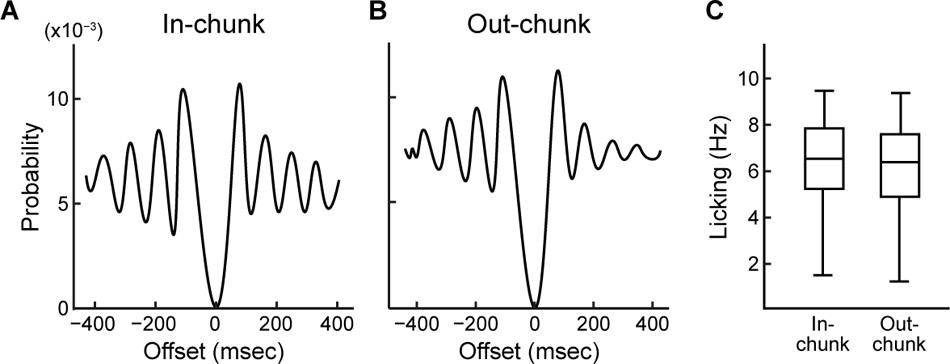
Licking auto-correlogram and frequency of in-chunk and out-chunk licks. A. Auto-correlogram of licking during in-chunk (left) and out-chunk (right) periods. B. Comparison of licking frequency during in-chunk (left) and out-chunk (right) periods for all chunks from all session. The median and interquartile range are shown in boxplots.

## STAR METHODS

### RESOURCE AVAILABILITY

#### Lead Contact

Further information and requests for resources and reagents should be directed to and will be fulfilled by the lead contact, Takashi Kitsukawa

#### Materials availability

This study did not generate unique reagent.

#### Data and code availability

- All data reported in this paper will be shared by the lead contact upon request.
- This paper does not report original code. All custom Python scripts will be available upon request.
- Any additional information required to reanalyze the data reported in this paper is available from the lead contact upon request.

## METHOD DETAILS

### Step-wheel

Detailed descriptions of the step-wheel can be found elsewhere (Kitsukawa et al., 2011; Nakamura et al., 2017). Briefly, the step-wheel is a motor-driven running Ferris-like wheel with series of pegs for the stepping surface serving as footholds for mice. The placement of pegs was changeable. We used ‘Regular’ and ‘Complex’ peg patters, designed as follows: In the regular peg-pattern (Regular), the pegs on the two sides were alternately spaced at constant inter-peg distances. In the complex peg-pattern (Complex), the pegs were arranged pseudo-randomly (Fig. 1C, see Kitsukawa et al., 2011 for details). Both peg patterns consisted of 24 pegs, 12 on each side, and a peg pattern was repeated twice in one rotation of the wheel (Fig. 1D). The passage through the entire peg pattern was counted as one trial. Thus, one rotation consists of two trials.

A spout was placed between the left and right side of pegs, from which mice could drink water as a reward if it kept pace with the turning wheel. Thus, the mice were required to run on the pegs at the same speed as the rotational speed of the wheel. Voltage sensors were installed on each peg and on the spout for detection of touches and licks (Kitsukawa et al., 2011). We have confirmed that body parts other than the paws (e.g., tail and head) were never detected by the voltage sensors. An infrared sensor detected a mouse close to the spout (Fig. 1B). All analyses were performed on data recorded while this infrared sensor was cut by the mouse.

### Mice

ICR mice (male, 10–20 weeks old, weighing 30–40 g) were purchased from SLC Japan (Hamamatsu, Japan). All procedures were approved by the Committee on Animal Care of the Massachusetts Institute of Technology and the Osaka University Animal Experiments Committee, and were performed in accordance with the National Research Council’s Guide for the Care and Use of Laboratory Animals.

The mice had free access to dry pellet food. During pre-training and training sessions, the mice were allowed to drink water principally in the step-wheel during and after the training session. Additional water was given in their home cages as needed to reach 3 ml/day. They were given free access to water for an entire day every 1–2 weeks. Their health condition, including weight, body temperature and fur condition, was closely monitored during water restriction.

Water was given ad lib if their weight fell to less than 80% of its initial level, or if any abnormal health condition was observed.

### Pre-training and training

To familiarize mice to running in the step-wheel while drinking water, we pre-trained them on the regular peg-pattern for 2–3 weeks. Pre-training continued until mice became able to run at the speed of 12.5 cm/sec. Training on the complex peg-pattern version of the task started on the next day after the completion of the pre-training (Fig. 1A).

Mice were trained in one session per day and performed 60–70 trials (1 rotation was defined as 2 trials) of the complex peg-pattern at a speed of 12.5 cm/sec. The training period was 9 days. Histograms of touch frequency (illustrated under the raster plots in Fig. 1E) were obtained by counting touch events (peg-touch) in every 5-msec bin and by smoothing over 10 bins.

### CV of inter-touch interval

All data analyses were conducted using custom Matlab (Mathworks, MA) programs. The touch intervals for all peg pairs (inter-touch interval) were calculated using the peg-touch data recorded. The CV (coefficient of variation) of the inter-touch interval in a session was calculated for each peg and shown in heat maps. The inter-touch CV of a target Peg_(N)_ was defined as the CV of the inter-touch intervals between the previous peg (Peg_(N-1)_) to the target peg (Peg_(N)_) calculated throughout a session. Heat maps were drawn by using the inter-touch CV calculated for each peg. In a given heat map, pegs with large inter-touch CV are drawn in darker gray, and pegs with small inter-touch CV are drawn in lighter gray (see Fig. 2A). The heat maps were normalized by setting the mean value to 1.

There were 24 pegs in the complex peg-pattern, but mice usually used only 18–22 pegs, and the number and locations of these pegs varied by mouse and by session. If the number of touches for a peg was less than one-third of the number of trials in the session, that peg was excluded from the analysis, and the inter-touch CV was calculated only among the remaining pegs. The excluded pegs were not used in the later analysis. All subsequent principal analyses were performed on data collected during the training period and not in the pre-training period, because no notable variations in inter-touch CVs were found during regular peg-pattern running. This stability was attributed to the even and alternatively spaced structure of the regular peg-pattern. Thus, the principal analyses focused exclusively on the training period, using the data collected as mice learned top ran on the complex peg-pattern.

### Definition of a chunk

To determine the boundaries of a chunk, we searched for pegs at which the inter-touch CV jumped up at both ends of the low-CV segment. Initially, the pegs at which the low-CV segment started or ended (start-peg or end-peg, respectively) were determined using the inter-touch CV at each peg in the following way:

1. To find a start-peg, the inter-touch CV of each peg was examined one by one in the opposite direction of the peg number (from 12 to 1). If the inter-touch CV was more than 1.3 times the value of the previous peg, the inter-touch CV was considered to have jumped up, and the last peg before the jump up was determined to be a putative start-peg.
2. To find the end-peg, the inter-touch CV of a peg was checked in the order of the peg number. If the value was 1.35 times or more than the previous peg, it was considered that a jump up has occurred, and the last peg before the jump up occurred was identified as a putative end-peg.
3. The span of pegs between the putative start-peg and end-peg identified in steps (1) and (2) was defined as the putative low-CV segment. If all the inter-touch CV of pegs in that segment were lower than the standard deviations of the inter-touch CV of all pegs in the peg pattern and that of the start-peg or end-peg, the segment was defined as a low-CV segment.
4. When the length of the low-CV segment was more than 4 pegs in a row, the sequence was identified as a chunk.

The thresholds for significant jump-ups and jump-downs of the inter-touch CV were both determined from the distributions of jump-ups and jump-downs calculated across all sessions in all mice. As a population deviating from the central distribution was identified, the upper ten percentile of jump-ups (and downs) was selected as the threshold that effectively differentiates between the two groups (Supplementary Fig.1). Consequently, the thresholds were determined to be 1.35 and 1.3, respectively, for jump-ups and jump-downs.

### CV analyses inside and outside of chunk

The start-pegs and the end-pegs belonged to the chunk in all analyses. The pegs inside the chunk were noted as in-chunk pegs in some of the following analyses, and all pegs not included in the in-chunk were assumed to be out-chunk. When showing the changes of the inter-touch CV, the inter-touch CV for the pegs defined as all pegs, in-chunk pegs, and out-chunk pegs were averaged for each session (see Fig. 2B). A one-factor ANOVA followed by Tukey-Kramer’s multiple comparison test was performed to test the amount of change in the transition of CV. The CV ratio in each session was calculated as (mean out-chunk CV) / (mean in-chunk CV). A one-factor ANOVA followed by Tukey Kramer’s multiple comparison test was performed to test the change of the amount of the CV ratio throughout the session. When the adjusted p-values were less than 0.05, the difference was considered significant.

### Similarity analysis of chunks

To compare quantitatively the structure of chunks across different sessions, a chunk vector was used, which was defined as a vector of 24 components corresponding to pegs that indicate the position of the chunk. Specifically, the component of the chunk vector was 1 if a peg belonged to a chunk and 0 if a peg did not belong to any chunk. To quantify the similarity of chunk structure between sessions, we calculated the cosine similarity between the chunk vectors. The cosine similarities were calculated across all sessions in all mice and shown as a heat map (see Fig. 3A).

Chunk vectors were arranged in a session order from top to bottom for each mouse to make a chunk binary map (see Supplementary Fig. 2). The pegs that were excluded from the analysis due to the small number of peg touches were complemented in the following way. An excluded peg was defined as an in-chunk peg only if two adjacent pegs on both sides were in-chunk pegs. Otherwise, any excluded pegs were defined as out-chunk pegs.

To compare the similarity of chunk structures within an individual mouse (intra-individual), the dot product of chunk vectors in different sessions of the same mouse was calculated. The similarity between individual mice (inter-individual), after enumerating the pair of sessions between different mice, was evaluated by randomly choosing the same number of pairs as the number of intra-individual session pairs and by calculating the similarity between the sessions.

A one-factor ANOVA followed by Tukey-Kramer’s multiple comparisons was performed to test the difference in chunk similarity between intra- and inter-individual. When the adjusted p-values were less than 0.05, the difference was considered significant.

### Indices for interval and phase

The interval-shift and phase-shift were used as indicators of motor coordination in repetitive steps (Fig. 4A, B). We define “step” and “cycle” before defining interval and phase, and then define the interval-shift and phase-shift. A *step* was defined as the time spent between the forelimb touching a peg and the other forelimb touching the next peg on the other side. A *cycle* was defined as the time taken for a forelimb to touch a peg and then for the same forelimb to touch on the subsequent peg of the same side (Fig. 4A). An *interval* was the time required for one cycle, and an *interval-shift* was defined as the time difference in intervals between two consecutive cycles. The ratio of the time required for one step within one cycle was defined as *phase*, and the difference of phases between two consecutive cycles was defined as *phase-shift*.

Let and be the time when the mouse touches the i-th (i = 1 to 12) pegs on, respectively, the left and right sides, and the interval-shift before and after the target peg was calculated as:

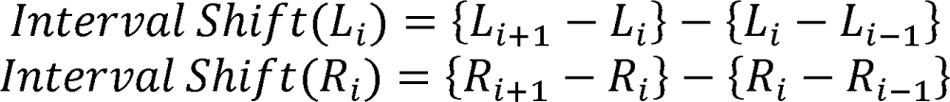

The interval-shift was calculated separately for the left and right trial by trial, then averaged over all trials in a session. The interval-shift of the opposite side to the start-peg and end-peg were aligned to the peg just before the start-peg or end-peg on the opposite side. The phase-shift of the target peg was calculated as:

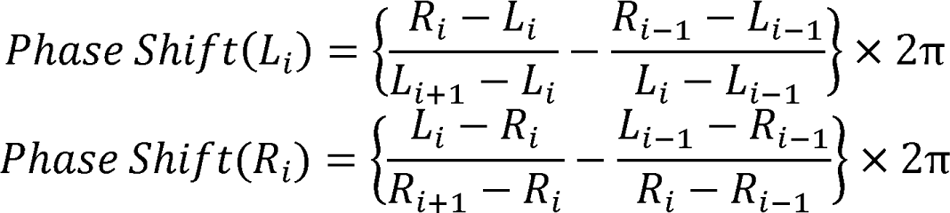

The phase-shifts were calculated trial by trial and were averaged across all trials in the session. This index shows how much the position of the opposite limb (phase) has changed before and after the target peg.

To analyze the distribution of interval, interval-shift, phase, and phase-shift, all the data from all sessions were used. The histogram of intervals (see Fig. 5A) was obtained by counting the number of intervals in each 10-msec bins and smoothed over 100 msec using sliding windows.

The histogram of the interval-shift (see Fig. 5B) was obtained by counting the number of interval-shifts in each 2-msec bins and smoothed over 20 msec using sliding windows. The histogram of phase (see Fig. 6A) was obtained by counting the number of phases in each 0.04 π (rad) bins and smoothed over 0.4 π using sliding windows. The histogram of the phase-shift (see Fig. 6B) was obtained by counting^π^ the number of phase-shifts in each 0.02 π bins and smoothed over 0.2 π using sliding windows. After the histograms of each variable were obtained, those were converted into the probability distribution.

### Comparison of interval-shift and phase-shift between in-chunk and out-chunk steps

For the comparison between in-chunk and out-chunk values for the interval-shift or the phase-shift, we used the data after the 4th session, in which chunk locations had become stable.

Changes in the mean interval-shift and phase-shift between in-chunk and out-chunk steps were tested by two-tailed t-tests. A one-factor ANOVA followed by Tukey-Kramer’s multiple comparisons was performed to test the changes in the interval- and phase-shift around start-pegs and end-pegs. When the adjusted p-values were less than 0.05, the difference was considered significant. For the per-mouse analyses, a one-factor ANOVA followed by Tukey-Kramer’s multiple comparisons was performed on the data from each mouse. When the adjusted p-values were less than 0.05, the difference was considered significant.

For the comparison of the phase-shift between chunk boundary region and other regions, we defined the boundary region as the region including start-peg and end-peg and one peg before and after such pegs (total of 6 pegs, three at the start and three at the end). All pegs except the boundary-region pegs were defined as belonging to the non-boundary region of the chunk. A two-tailed t-test was performed to test the difference in phase-shift between the boundary and the non-boundary regions. P-values less than 0.05 were considered significant.

### Licking analysis

Histograms of licking intervals were obtained by counting consecutive intervals in 2-msec bins and smoothing over 10 bins using sliding windows. For auto-correlograms, the licking events detected during the 500-msec windows before and after each licking were counted in 5-msec bins smoothed over 10 bins with sliding windows. Finally, the histograms were converted into probability distributions.

Auto-correlograms of licking events were constructed separately for the in-chunk and out-chunk regions with the same method described above (see Supplementary Fig. 6A). The period corresponding to in-chunk for licking analysis was determined as the time from touching start-peg to touching end-peg, and all other periods were determined as the time corresponding to out-chunk. The average number of licks for in-chunk and out-chunk was calculated for each mouse, and the means and SEMs were shown for all mice (see Supplementary Fig. 6B).

Throughout a given session, the number of licks that occurred in in-chunk and out-chunk periods was counted. Then, counts were divided by the total time that mice spent in each region to calculate the average number of licks per second.

For cross-correlograms between the licks and the peg touches, licks that occurred in the 300-msec periods before and after each peg touch were counted. A cross-correlogram was obtained by counting licks in each consecutive 5-msec bin (see Fig. 7C). A periodicity index of licking around each peg touch was calculated as the maximum value of the power in the 8–10 Hz range in the spectrogram, which corresponded to the frequency band of licking. A one-factor ANOVA followed by Tukey-Kramer’s multiple comparison test was performed to test the changes in the periodicity indices around start-pegs and end-pegs; adjusted p-values less than 0.05 were considered significant. For the per-mouse analyses, a one-factor ANOVA followed by Tukey-Kramer’s multiple comparisons was performed on the data of each mouse. When the adjusted p-values were less than 0.05, the difference was considered significant.

